# CUT&RUNTools: a flexible pipeline for CUT&RUN processing and footprint analysis

**DOI:** 10.1101/529081

**Authors:** Qian Zhu, Nan Liu, Stuart H. Orkin, Guo-Cheng Yuan

## Abstract

We introduce CUT&RUNTools as a flexible, general pipeline for facilitating the identification of chromatin-associated protein binding and genomic footprinting analysis from antibody-targeted CUT&RUN primary cleavage data. CUT&RUNTools extracts endonuclease cut site information from sequences of short read fragments and produces single-locus binding estimates, aggregate motif footprints, and informative visualizations to support the high-resolution mapping capability of CUT&RUN. CUT&RUNTools is available at https://bitbucket.org/qzhudfci/cutruntools/.

---

Mapping the occupancy of DNA-associated proteins, including transcription factors (TF) and histones, is central to determining cellular regulatory circuits. Conventional ChIP-sequencing (ChIP-seq) relies on cross-linking of target proteins to DNA and physical fragmentation of chromatin^1^. In practice, epitope masking and insolubility of protein complexes may interfere with successful use of conventional ChIP-seq for some chromatin-associated proteins^2–4^. CUT&RUN is a recently described native endonuclease-based method based on binding of an antibody to a chromatin-associated protein *in situ*, and the recruitment of a protein A-micrococcal nuclease fusion (pA-MN) to antibody to efficiently cleave DNA surrounding binding sites^5^. The CUT&RUN method has been successfully applied to a range of TFs in yeast^5,6^ and mammalian cells^7,8^. The procedure achieves higher resolution mapping of protein binding since endonuclease digestion generates shorter fragments than physical fragmentation. In our experience, existing tools to analyze such data proved inadequate due to the lack of an end-to-end computational pipeline specifically tailored to this technology. Therefore, we have developed a new pipeline, designated CUT&RUNTools, that streamlines the processing, usage, and visualization of data generated by CUT&RUN (Fig 1a).

**Figure 1:**
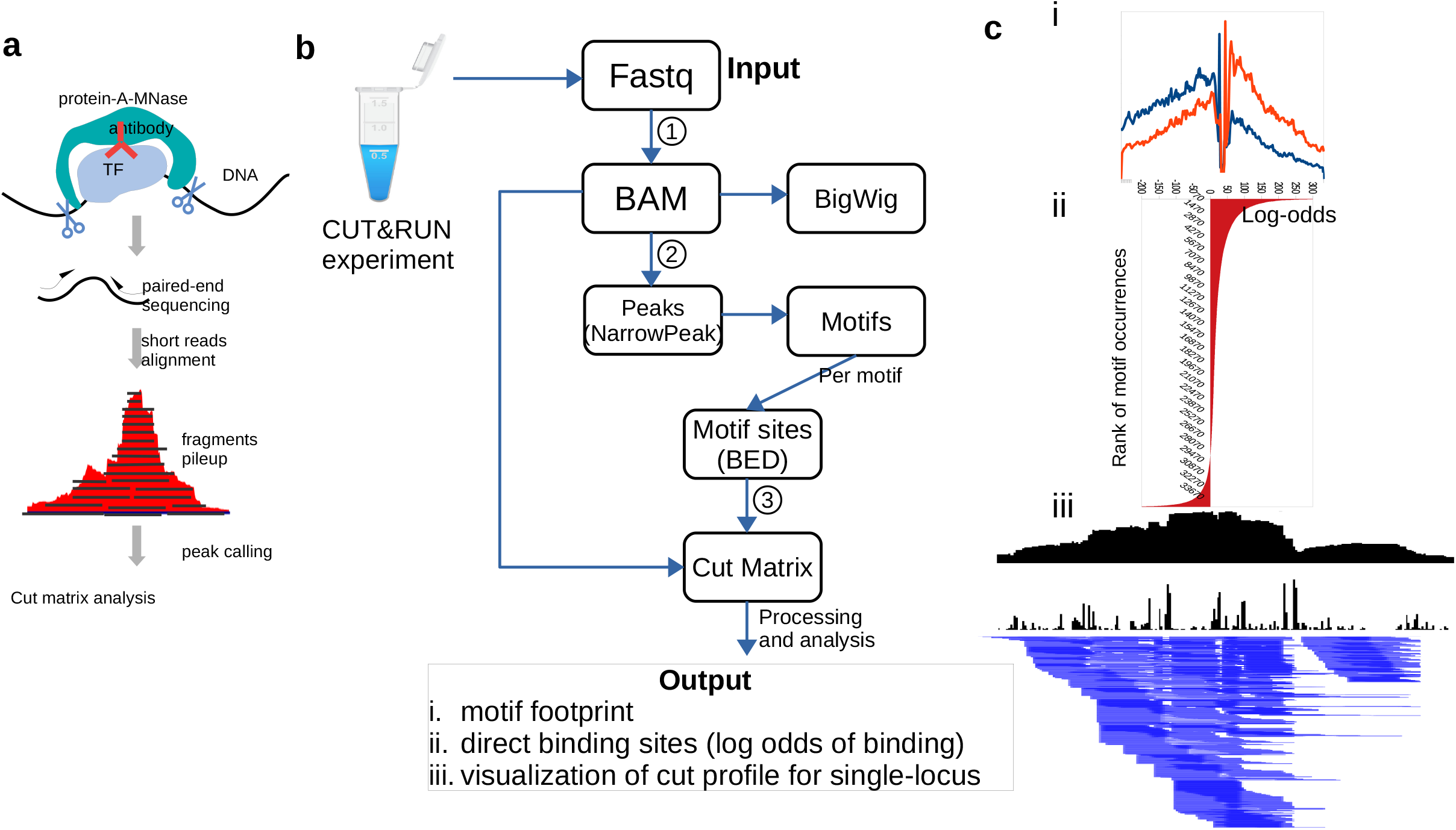
a. Schematic of CUT&RUN. pA-MN is recruited to TF-bound antibody and cleaves around TF binding site, liberating DNA fragments for sequencing. Subsequent steps require a specially designed computational pipeline to extract maximal information from the data. b. Overview of CUT&RUNTools. Step 1: input paired-end raw reads are aligned to the reference genome with special care for short read trimming and alignment. Step 2: peaks are called based on fragment pileup. A fixed window around the summit of each peak is used to perform *de novo* motif finding. Step 3: the cut matrix is calculated for each motif of interest and used to generate the three outputs: i) motif footprint, ii) direct binding site identification, and iii) visualization. c. The output of CUT&RUNTools at the chr3:98302650-950 region as an example.

CUT&RUNTools takes paired-end sequencing read FASTQ files as the input, and performs a set of analytical steps: trimming of adapter sequences, alignment to the reference genome, peak calling, extraction of cut matrix at single-nucleotide resolution, *de novo* motif searching, motif footprinting analysis to determine the unique footprint, and identification of individual binding sites (Fig. 1b). The outputs of the pipeline (Fig 1c) are: 1) an aggregate footprint capturing the characteristics of chromatin-associated protein binding (Fig 1c, i), 2) binding log odds values for individual motif sites informative for direct binding sites identification (Fig 1c, ii), and 3) visualization of a cut frequency profile at nucleotide resolution (Fig 1c, iii).

Specifically, CUT&RUNTools performs alignment with special attention to short-read trimming and read alignment (Fig 1b, **step 1**)(Methods). Due to the predominance of short fragments (25-50 bp) generated by CUT&RUN, the typical settings in read trimming and sequence alignment does not perform well. We introduce a two-step read trimming process to improve the quality. First, the sequencing data are processed with Trimmomatic^9^, a commonly used template-based trimmer. Next, a second trimming step was included to remove any remaining adapter overhang sequences not removed due to fragment read-through. CUT&RUNTools further adjusts the default alignment settings by turning on dovetail alignment^10^, designed to accept alignments for paired-end reads when there is a large degree of overlap between two mates of each pair. Together, this improved trimming and alignment procedure increased the alignment percentage from 68% to 98% compared to a setting with no trimming and alignment adjustments (**Supplementary Table 1**). With the reads aligned, CUT&RUNTools employs MACS^11^ to perform peak calling based on the coverage profile, followed by *de novo* motif searching within peak regions with MEME suite^12^ (Fig 1b, **step 2**).

An important element of CUT&RUN analysis is the estimation of cut sites, which enables more accurate positioning of binding locations than simple peak calling. The cut sites derive from the two ends of individual DNA fragments generated upon cutting of chromatin by the pA-MN fusion recruited to the antibody binding sites. Regions of lower cut frequency tend to be protected due to chromatin-associated protein binding, whereas flanking regions without binding display higher cut frequencies (**Supplementary Fig 1a**). CUT&RUNTools accurately tabulates the frequency with which cleavage is observed at each base pair (Methods).

Using the cut matrix, footprinting analysis^13,14^ is then applied to identify high-resolution occupancy of sequence-specific binding factors such as TFs. To detect footprints from CUT&RUN data, CUT&RUNTools first generates an aggregated cut frequency profile based on all +/- 100bp regions extending from each peak-embedded motif site. Then, CUT&RUNTools estimates a probabilistic bimodal clustering model ^15^, and assigns a binding probability score, expressed as log-odds, to each motif occurrence based on the model. The score, log-odds of TF being bound, is a reflection of how well the cuts at each motif occurrence matches the aggregate footprint pattern. By ranking sites by the score, CUT&RUNTools generates a rank-ordered list of likely direct binding sites. For histone proteins and epigenetic factors that do not have clear sequence specificity, the motif and footprint analyses may be skipped.

We illustrate the functionality of CUT&RUNTools through analysis of CUT&RUN data acquired for GATA1, a master regulator in erythroid lineage cells^16^. We performed CUT&RUN using GATA1 antibody in primary human stem/progenitor CD34+ cells after 7 days of erythroid differentiation (Fig 2). Results were compared initially to published GATA1 ChIP-seq data for cells under the same conditions^7^. Peaks identified in CUT&RUN align very well with ChIP-seq at over 35,000 GATA1 sites across the genome (Fig 2a). Furthermore, the pileup signal in CUT&RUN is more enriched in a narrower window in the peak center than ChIP-seq (50bp vs. >150bp), reflecting higher resolution (Fig 2b). As expected, CUT&RUNTools correctly identified the HGATAA GATA1 recognition motif *de novo* (E=1e-200). Next, we performed GATA1 footprinting using the cut-matrix generated on the HGATAA motif by CUT&RUNTools and the surrounding 150bp regions for all 35,000 sites in peak regions (Fig 2c). Indication of protection in the motif core was particularly strong (Fig 2c, e). Based on the estimated log odds scores (Fig 2d) (**Supplementary Table 2**), CUT&RUNTools identified 25,900 of the 35,000 motif sites as direct binding sites. Comparison with literature data validates these estimates (**Supplementary Fig 2**) (in addition, a systematic comparison with ChIP-seq is shown in Fig 2a). Of note, a stereotypical, center-depleted cutting pattern is identifiable not only from the average profile but also at individual motif sites.

**Figure 2:**
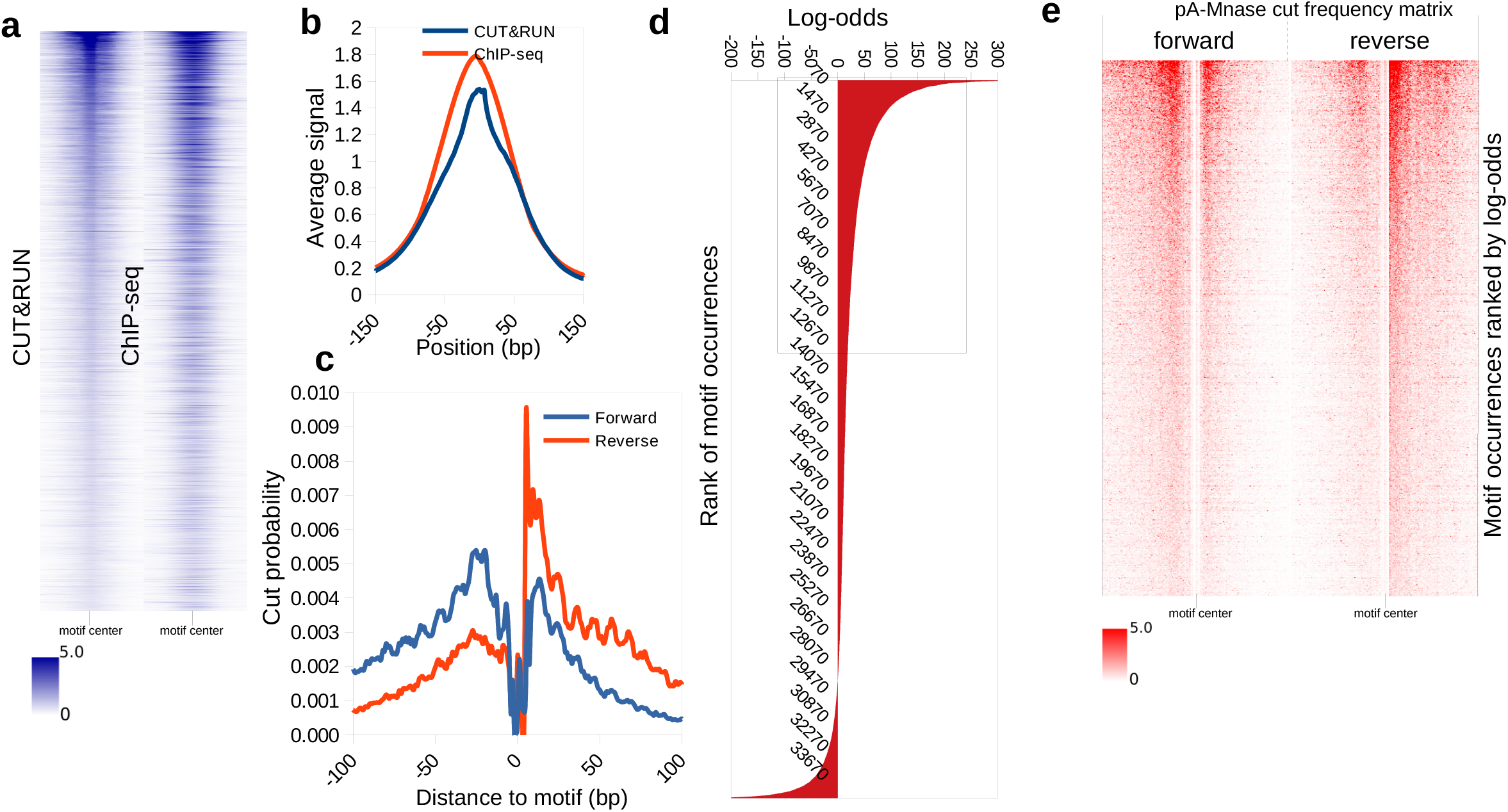
a. GATA1 CUT&RUN and ChIP-seq comparison. GATA1 motif is scanned across CUT&RUN and ChIP-seq peaks. The signal pile-up of −150bp to +150bp region surrounding each motif site is plotted. b. CUT&RUN signal is enriched in a narrower window than ChIP-seq, consistent with higher resolution of the CUT&RUN method. c. CUT&RUN footprint for the HGATAA motif. Enzyme cut protection is noted in motif core, and deprotection in the flanking regions. d. The distribution of log-odds score for genome-wide HGATAA motif sites. A threshold value of 5 is used to determine direct binding sites. e. Strand-specific cut frequency profile at individual HGTAA motif sites, illustrated as a heatmap.

In addition, *de novo* analysis of GATA1 CUT&RUN returned several additional motifs that may correspond to co-factors (**Supplementary Data 1**). These co-factor motifs (also termed secondary) can be distinguished via an asymmetrical motif footprinting profile (**Supplementary Fig 4**), in contrast to the symmetrical profile of the primary HGATAA motif (Fig 2c). We use a ‘footprint symmetry score’ (FSS) to discriminate primary from secondary motif footprints (Methods, **Supplementary Fig 3a,b**) (**Supplementary Table 3**). HGATAA has the highest FSS score (**Supplementary Fig 3c**). Identified co-factor motifs GCCCCGCCCTC, CMCDCCC, RTGASTCA (**Supplementary Fig 3d**) correspond to SP1, KLF1 and NFE2 TFs, respectively, which are known to cooperate with GATA1^17^. Each displays a noticeably higher rate of descent on one side of motif than the other (**Supplementary Fig 4**).

Importantly, *de novo* analysis also identified an extended motif for co-binding of GATA1 and TAL1^18^. GATA1 forms a multiprotein complex with TAL1 along with LMO2 and Ldb1^18,19^. The GATA1-TAL1 complex recognizes HGATAA and a half E-box (TAL1) separated by a gap of 10 nucleotides^20^. Despite the length of this motif, CUT&RUNTools displays a strong footprint shape for the extended motif, with high FSS indicating that it is primary, and lending support to the known complex binding model (**Supplementary Fig 5**). The motif footprinting is consistent between *de novo* and known GATA1-TAL1 motifs (**Supplementary Fig 5**). Therefore, in cases where the recognition sequence of TF is not known in advance, *de novo* analysis in combination of genomic footprinting should be helpful in establishing the primary motif for the TF and characterizing motifs for co-factors.

Tools are available for ATAC-seq and DNase-seq data analysis that enumerate cut frequencies and construct cut matrices^21,22^. In practice, however, we found that direct application of such tools to analyze CUT&RUN data leads to incorrect cut positions (**Supplementary Figs 6, 7**). One reason is that the two ends of paired reads do not each indicate an end of a fragment (**Supplementary Fig 8**), while only the 5’-end of a read does, making the accounting of cut positions error-prone. Another important difference is that Tn5 transposase in ATAC-seq leaves a 4bp overhang in sequenced fragments^23^, whereas pA-MN enzyme in CUT&RUN cleaves surrounding the location of binding sites with no overhang. Specific adjustments are thus required and have been made in the enumeration of cut matrix to take into account this feature of CUT&RUN (Methods). Recognizing these differences, we provide an option to tune the cut site offset to make CUT&RUNTools applicable for both CUT&RUN and ATAC-seq footprinting analyses (**Supplementary Fig 9**) and in doing so allow flexibility of experiment type.

Finally, CUT&RUNTools includes several quality control metrics, including fragment size distribution, reads duplication rate, library size, adapter content percentage, alignment percentage to assist users in quality control evaluation of CUT&RUN experiments (Methods). By further using the number of peaks and the enrichment of the expected motif, users can evaluate overall success of experiments and validate a given antibody. Additionally, CUT&RUNTools generates publication-quality visualizations to aid biologists in interpreting cleavage data and to substantiate evidence of binding (Fig 3). The cut frequency track, for example, displays the number of cuts at each nucleotide position within a specified genomic range. A broad-level visualization (300bp) (Fig 3a) highlights the location of motif and other footprints within the region. At 100bp resolution (Fig 3b), a genomic sequence view is enabled and the exact locations of cleavage can be seen. These visualizations can be executed simply through user-friendly commands. CUT&RUNTools supports SLURM^24^-based cluster environment, and permits simple specification of inputs/outputs, tools, and resource-related parameters through a JSON-formatted configuration file. A detailed usage manual is provided online regarding these settings and various usage scenarios.

**Figure 3:**
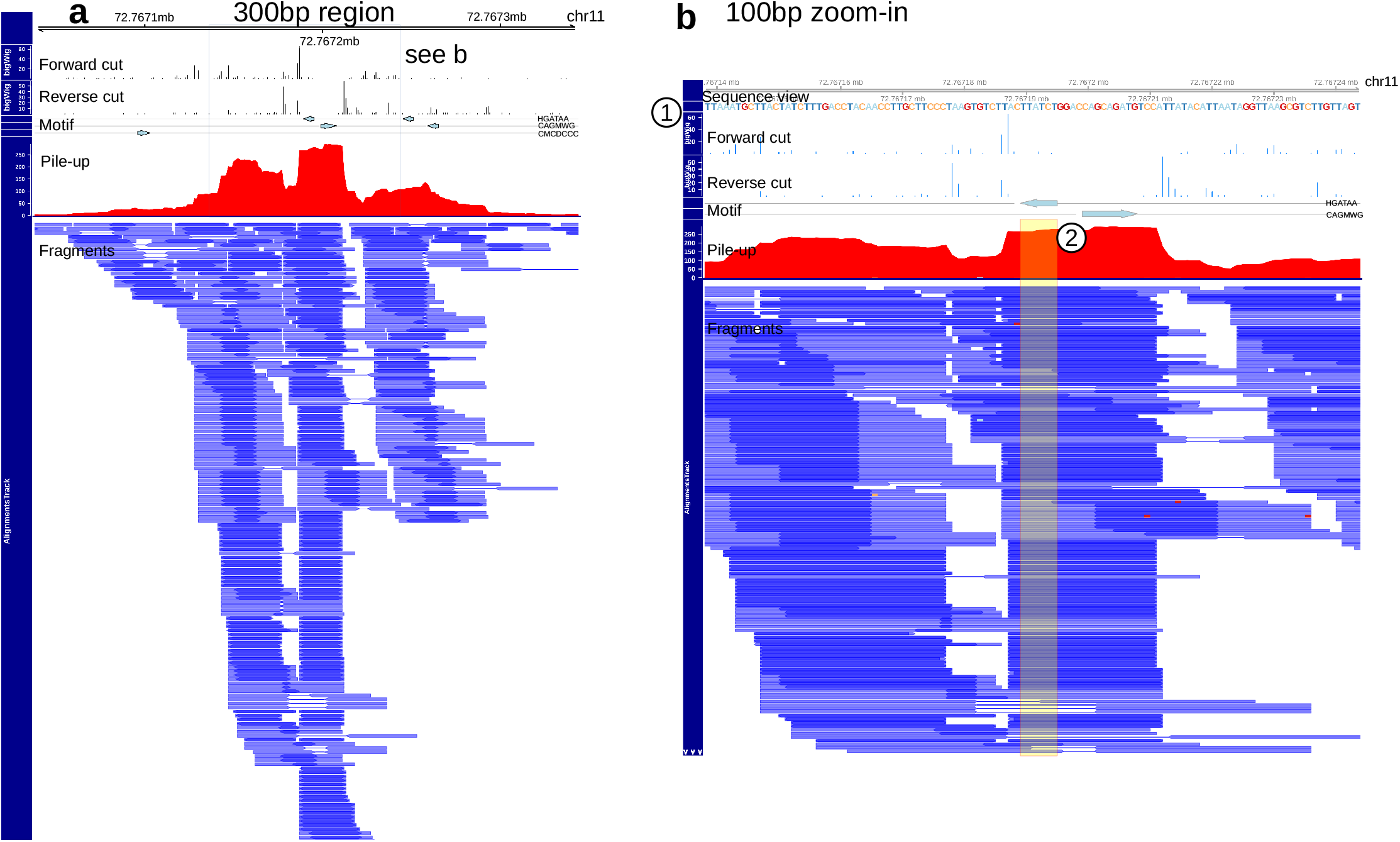
a. CUT&RUNTools visualization of an example region chr11:72767100-72767300. The top two tracks show the strand-specific cut frequency profiles. Third track is signal pile-up plot. Fourth track is the fragment plot, showing the location, start and end positions of each DNA fragment. Forward cut frequency refers to the end of R1 mate and reverse refers to the end of R2 mate, where the designation of R1, R2 is based on whether the mate alignment contains the motif, or the motif’s reverse-complement. b. A zoom-in view of the same region as a. The view contains an additional sequence view (1) and highlighting of HGATAA motif (2).

In summary, CUT&RUNTools provides a means of directly detecting TF binding through assessment of the protection of TF-bound DNA from enzyme cleavages, and we illustrate that such cut site information is valuable in the accurate identification of TF binding sites. Thus, CUT&RUNTools should enable biologists to realize advantages of cleavage data provided by CUT&RUN, and make high-quality footprinting analysis accessible to a broad audience.

Supplementary Figure 1:

a. Illustrative example of a cut frequency matrix. Rows: motif sites. Columns: (−100bp, motif, +100bp) region. Cuts are tabulated at each position of the motif site. Darkness of red indicates more cuts.

b. Steps to construct an aggregate motif footprint. First, all peaks are scanned for motif sequence. Then, for each motif occurrence, we obtain cut frequency on (−100bp, +100bp) region centered on motif. Next, CENTIPEDE computes the posterior probability of cutting at each position of the motif site using an iterative procedure based on the spatial distribution of reads around motif. The output is the posterior probability of cutting per position, a number weighted from all motif sites according to each site’s overall probability value.

Supplementary Figure 2:

Validation of two GATA1 sites at the CPOX gene locus. GATA1 binding at these sites is confirmed based on a previous report ^25^. CUT&RUNTools predicts binding with highly significant log odds of 315 and 115 respectively, which are ranked in the top 0.1% and 2% of all sites. A zoom-in view is provided on each site to show 1) ChIP-seq profile, 2) CUT&RUN coverage profile, 3) CUT&RUN cut frequency profile. Cut frequency profile shows a series of footprints that may correspond to GATA1 and other transcription factors.

Supplementary Figure 3:

Footprint symmetry analysis reveals the primary and secondary motifs.

a. Symmetry analysis calculates the ascent, and descent rates for the two sides of footprint via fitting an exponential decay curve to each part.

b. An example of exponential decay curve that is fit on real data.

c. A perfect symmetry indicates a primary motif (HGATAA).

d. A substantial difference between ascent, and descent rates would indicate a motif footprint with a asymmetric footprint shape, as shown by CMCDCCC, MGGAAR, RTGASTCA secondary motifs. Asymmetry is quantified by the Footprint Symmetry Score (FSS).

Supplementary Figure 4:

Examples of secondary motifs detected from GATA1 CUT&RUN experiment. Secondary motifs in general have asymmetrical footprint shapes and usually indicate factors that co-occur with the primary antibody. From left to right, these motifs correspond to factors KLF1, ETS1, RUNX1, NFE2, and KLF1.

Supplementary Figure 5:

GATA1-TAL1 composite motif footprint (a) using the motif that is found *de novo*, and (b) using the motif that is from JASPAR motif database.

Supplementary Figure 6:

Comparisons of HGATAA motif footprint using the cut matrix enumerated by (a) CUT&RUNTools, (b) Atactk, and (c) CENTIPEDE.tutorial. Options b) and c) are unsuitable for CUT&RUN data as they are specifically designed for ATAC-seq and DNase-seq data respectively. They miscalculated the cut matrix. See Supplementary Fig 7 for numerical comparisons of the cut matrices.

Supplementary Figure 7:

a. Cut matrix comparison between CUT&RUNTools, CENTIPEDE.tutorial, and Atactk. Cut frequency is presented as a bar plot on HGATAA motif site that is located at **chr1:32510-32516, strand (−)**. Plot shows cuts in 206bp regions surrounding this site. The first 206bp indicates cuts on forward strand, and the next 206bp shows reverse strand. Insert (i) shows the zoom-in of first 20bp of CUT&RUNTools and Atactk, showing a 4bp shift between their profiles. CENTIPEDE.tutorial and Atactk both miscalculated cuts when applied to CUT&RUN data. Right-hand side shows the reason for the errors. Specifically, the cut matrix generated by CENTIPEDE.tutorial should be reversed since motif is on (−) strand, and Atactk’s matrix is off by 4bp.

a. Cut matrix comparison between CUT&RUNTools, CENTIPEDE.tutorial, Atactk. Cut frequency is shown for HGATAA motif located at **chr1:32592-32598, strand (+)**. Plot shows cuts in two 206bp regions surrounding motif (−100bp, motif, +100bp). First 206bp: forward strand. Next 206bp: reverse strand. Insert (i) shows the zoom-in of first 20bp of CUT&RUNTools and Atactk. Atactk contains errors in the estimated cuts. Right-hand side shows the reason for error.

Supplementary Figure 8:

Anatomy of DNA fragment with both mates of a pair (R1, R2) indicated. Three scenarios of a DNA fragment could occur: i) R1 and R2 has a gap (unsequenced region) in the middle. ii) R1 and R2 overlap partially. iii) R1 and R2 completely overlap. Ends of a read do not always correspond to a fragment end, due to unsequenced region.

Supplementary Figure 9:

Application of CUT&RUNTools on ATAC-seq data. Motif footprinting analysis was performed on two given motifs, GATA1, and CTCF on ATAC-seq HUDEP-2 cells. CUT&RUNTools was able to generate a motif footprint in each case. A cut site offset of 4bp was used.

Supplementary Figure 10:

a. With cut site offset set to 0, atactk-generated cut matrix creates a motif footprint that is close to the correct solution (see b), but is still incorrect because the forward and reverse strands are misaligned by 1bp (see insert).

b. The correct motif footprinting plot estimated by CUT&RUNTools.

## Methods

### CUT&RUN experiments

CUT&RUN experiments were carried out following the nuclei isolation version of protocol as described^5,7^. Nuclei from 2×10^6^ cells were isolated with NE buffer that consisted of 20mM HEPES-KOH pH 7.9, 10 mM KCl, 0.5 mM Spermidine, 0.1% Triton X-100, 20% Glycerol and 1x protease inhibitor cocktails. Nuclei were captured with BioMagPlus Concanavalin A and incubated with 2μg primary antibody (α-GATA1, ab11852, abcam) in 200 μL wash buffer (20mM HEPES-NaOH pH 7.5, 150 mM NaCl, 0.5mM Spermidine, 0.1% BSA and 1x protease inhibitor cocktails) for 2 hours. Then unbound antibody was washed away with 400 uL wash buffertwice. Then pA-MN was added at 1:1000 ratio to 200μL wash buffer and incubated for 1 hour. Nuclei were washed again and resuspended in 150μL wash buffer. CaCl_2_ was next added at a final concentration of 2mM to activate the enzyme. Reaction was carried out at 0°C and stopped by 150μL of 2X STOP buffer (200mM NaCl, 20mM EDTA, 50 ug/mL RNase A and 40μg/mL glycogen). Protein-DNA complex was released by centrifugation and digested by proteinase K at 50°C overnight, followed by DNA precipitation by ethanol. The pellet was washed with 70% ethanol and dissolved in 25 μL 0.1x TE (1 mM Tris-HCl ph 8.0, 0.1 mM EDTA).

#### CUT&RUN libaray preparation and sequencing

The NEBNext Ultra II DNA Library Prep Kit was used with modifications described previously^7^ which aims to preserve short DNA fragments (30-80bp). Briefly, 6 ng of CUT&RUN DNA were treated with endprep module at 20°C for 30 min, 50 °C for 1 hour to reduce the melting of short DNA. Ligation was performed by adding 5 pmol of NEB adapter and ligation mix, and incubated at 20 °C for 15 min. To clean up the reaction, add 1.75x volume of Agencourt AMPure XP beads (Beckman Coulter) to capture short ligation products. PCR amplification was performed for 12 cycles. The resulting libraries were purified with 1.2x volume of AMPure beads, then analyzed and quantified by Qubit and Tapestation. The detailed step by step protocol can be found at protocol.io (https://dx.doi.org/10.17504/protocols.io.wvgfe3w). Libraries with different indexes were pooled and Illumina paired-end sequencing was performed using Nextseq 500 platform with NextSeq 500/550 High Output Kit v2 (75 cycles) (2×42bp, 6bp index).

### CUT&RUNTools implementations

Broadly, CUT&RUNTools consists of trimming, alignment, peak calling, motif finding, and cut matrix generation, and motif footprinting steps. The pipeline incorporates specific changes to some of the steps to accommodate the short read, short fragment characteristics of CUT&RUN. Its cut matrix generation ensures an accurate accounting of cut positions for footprint analyses. These steps are described below.

#### Raw reads trimming and alignment

Short fragments are frequently encountered in CUT&RUN experiments due to the fine cutting by pA-MN enzyme. As a result, it is common to expect both mates of DNA fragment to overlap. When the fragment is shorter than the length of a read, then we can expect that adapter run-through will occur. It is thus critical to remove adapter sequences at the end of reads. To deal with issues caused by alignment of short fragments, we made two important modifications to the typical adapter trimming and alignment protocol:

1. An initial trimming was first performed with Trimmomatic^9^, with settings optimized to detect adapter contamination in short read sequences. Trimmomatic is a template-based trimmer. However, reads containing 6bp, or less, of adapters are not trimmed. Therefore, a separate tool Kseq was developed to trim up to 6bp adapters from the 3’ end of each read that was not effectively processed by Trimmomatic. Note that this trimming does not affect the cut site calculation, which counts only the 5’ end of sequences. After trimming, a minimum read length of 25bp was imposed, as reads smaller than this were hard to align accurately.
2. Dovetail alignment policy. Bowtie2^10^ aligns each mate of a pair separately and then discards any pairs that have been aligned inconsistently. Dovetail refers to the situation when mates extend past each other. In the default setting, these alignments are discarded. Dovetail is unusual but encountered in CUT&RUN experiments. The --dove-tail setting^10^ was enabled to flag this situation as normal or “concordant” instead of elimination of such reads.

#### Peak calling and motif finding

After alignment, fragments that were longer than 120bp were filtered away. Then MACS2 was applied with the default narrowPeak setting^11^. Afterward, sequences within 100bp from the summit of each peak were obtained, and any sequences containing a substantial amount of repeats (as reported by RepeatMask) were removed. These remaining sequences were next used to perform *de novo* motif searching using MEME^12^. The top 20 motifs were saved for subsequent analyses. FIMO (part of MEME suite^12^) was applied to enumerate all motif sites in peak regions.

Like other techniques, some fraction of sequenced read pairs appear as duplicates (i.e. with identical start and end positions between duplicates). However, it is argued that nuclease cleavage of chromatin by its stereotypical nature is influenced by conformation of chromatin and/or nuclease bias^26^, and shorter DNA fragments also increased the likelihood of identical reads that originated from different cells^27^. Thus, removing duplicates from CUT&RUN experiments should be dealt with caution if the library complexity is not too low (due to extremely low input and/or high PCR cycle numbers). Thus, the default action in CUT&RUNTools is to retain duplicate reads, and users can choose to remove duplicates at their own discretion. We recommend users to be aware of low complexity of libraries with high duplication rates, as these may indicate a poor quality preparation. Users may repeat peak calling analysis on both duplicate and duplicate-removed instances. By comparing peak number, motif enrichment, enrichment of expected motifs, and other quality metrics, users may decide whether it makes sense to use the duplicate version for subsequent analysis.

#### Cut matrix generation

For any motif of interest, its corresponding cut matrix was generated as follows. The rows of cut matrix are the motif sites. The columns are the individual nucleotides in the −100bp, motif, and +100bp regions. Cut matrix requires all motif sites to be in a consistent orientation. That is, if the motif occurrence is located on the minus strand in the reference genome, all the cut frequencies in that motif site are flipped, so that −100bp position from the old profile becomes the +100bp position in the new profile. By convention, a value at i-th nucleotide means the cut is situated just before i-th nucleotide. The cut matrix tabulates the frequency of fragments ending in each nucleotide.

To compute strand-specific cut matrix, the ends of DNA fragments that overlap with the motif were assigned to forward and reverse strand cut matrices as follows. For each fragment, define R1 and R2 as two mates. The ends of the fragment are the start of R1 (s1) and the end of R2 (e2). If a given motif occurrence appears on the positive strand of the reference genome, then s1 belongs to the “forward” strand cut and e_2_ belongs to the “reverse” strand cut. Otherwise if the motif occurrence is on the negative strand, then s1 belongs to the “reverse” strand cut and e2 belongs to the “forward” strand cut. Likewise, tabulation was repeated for all paired-reads and for all motif occurrences, each time separately for each strand.

#### Motif footprinting analysis

A motif footprint is a plot that shows the enzyme cleavages around the motif region, presumabiy due to the protection of TF-bound DNA. It is typically characterized by a low cut frequency (or low posterior probability of cut) in the motif core, and a high cut frequency in the motif flanking regions. Prior to footprint analysis, blacklisted regions were excluded from the the peak list. Any chromosome M peaks were also excluded. Next, CENTIPEDE^15^ was applied to fit a probabilistic bimodal clustering model on the strand-specific cut matrix data which has aligned and centered all motif containing regions. CENTIPEDE was run with default settings and specifying the length of the motif.

#### Footprint symmetry analysis for identification of primary and secondary motifs

CUT&RUNTools has built in a feature to determine whether a motif footprint is primary or secondary, based on a ‘footprint symmetry score’ (FSS) defined as follows. The footprint profile is first divided in the middle into two halves, and to capture shape information, each half is fitted by a exponential decay curve (of the form A_left_ exp(B_left_ * x) and A_right_ exp(B_right_) * x, repectively) (**Supplementary Fig 3**). The parameter B_left_ (and B_right_, respectively) reflects the ascent rate for the left arm (and the ‘descent rate’ for the right arm, respectively). The goodness of fit is quantified using the R-square statistic, represented by R^2^_left_ and R^2^_right_, respectively. The FSS score is defined as B_letf_ * R^2^_left_ + −1 * B_right_ * R^2^_right_. Intuitively, the FSS score meaures the rate of increase of cut probabilities in the footprint plot, as the position approaches the motif. This rate should match the respective rate of decrease of cut probabilities as the position is further away from center. A FSS score of >0.3 and a small difference between B_left_ and −1*B_right_ indicates symmetry of motif footprint. Such a motif is designated primary.

#### Determining direct binding sites

The criteria we used for direct binding sites were following: 1) the site must contain a primary motif; 2) the site must fall within a CUT&RUN peak; 3) the site must have a high binding log-odds, which assesses the compatibility of the cut frequencies at the site with the binding model. Binding log odds, estimated by CENTIPEDE, is defined as log(p/(1-p)) where p is the overall posterior probability of binding at each site. The posterior probability for bound case (p) is estimated from a multinomial distribution and uses information from the spatial distribution of reads around the motif:

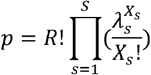

Where R is the total of reads in the region (modeled with a negative binomial distribution), s is a position index in the motif, *λ_s_* is the per position posterior probability of cutting, X_s_ is the per position number of reads. In the null case (no binding), *λ_s_* is equal to 1/S or uniform. Because posterior log odds log(p/(1-p)) is a likelihood ratio, its estimation can use a shorter derived form for simpler numerical computation (see CENTIPEDE^15^). Running CENTIPEDE on a primary motif would satisfy the first two of three criteria already, since footprinting is performed on CUT&RUN peak regions only. Based on the CENTIPEDE result, we set a stringent cut off of log-odds >5 to obtain direct binding sites for the motif.

#### Pipeline implementations

CUT&RUNTools was implemented using Python, R, and BASH scripts. Visualizations of motif footprints were implemented using matplotlib library in Python. Visualization of single-locus cut profile was implemented using the Gviz R package^28^. Integration of next-gen sequencing tools was achieved using Python and BASH scripts. Configuration of pipeline, including inputs/outputs, prerequisite paths, is specified by a JSON formatted file. CUT&RUNTools works under the SLURM^24^ job submission environment. A usage manual is provided online at the repository link: https://bitbucket.org/qzhudfci/cutruntools.

#### Comparison with other cut matrix tools

There are two currently available tools for enumerating cut matrices from enzyme cleavage data. One is Atactk, designed for ATAC-seq data, and the other is CENTIPEDE.tutorial, targeted towards DNase-seq. These tools were each applied to CUT&RUN data for the purpose of showing the advantage of CUT&RUNTools. Make-cut-matrix tool from the Atactk package^21^ v0.1.5 was downloaded from (https://github.com/ParkerLab/atactk) and the CENTIPEDE.tutorial package v1.0 was downloaded from https://github.com/slowkow/CENTIPEDE.tutorial. Make-cut-matrix was run with default settings on GATA1 CUT&RUN data, using HGATAA as motif. The centipede_data() function of CENTIPEDE.tutorial package was used to generate cut matrix with default parameters. To evaluate the quality of cut matrix generated by these tools, CENTIPEDE motif footprinting was performed on the generated cut matrices, and the quality of motif footprint plot was inspected for differences. Two loci was selected to more specifically compare the cut frequency profile estimated by these tools and CUT&RUNTools, and illustrate their differences.

#### Cut matrix implementation in CUT&RUNTools

To make sure that cut matrix is accurately estimated for CUT&RUN data, CUT&RUNTools adapts the following changes starting with the make-cut-matrix implementation. Adjustments are written in the form of a ‘patch’, which is available in the pipeline. First, the default setting of 4bp cut site offset was removed as it was usually required for ATAC-seq data (due to Tn5 transposase imposing a 4bp overhang on the sequences^23^). CUT&RUN cuts approximately at the TF binding site, so no cut site offset is required (offset = 0). Second, the position of reverse strand cut site is noted to be shifted by 1bp even after setting cut site offset to be 0 (**Supplementary Fig 10a**). This shift has been a remnant feature of ATAC-seq where forward strand has a cut offset of 4bp while the reverse strand has a cut offset of 5bp. So an adjustment of cut position has been further made to correct this behavior (**Supplementary Fig 10b**). With both of these changes adapted, the cut matrix was independently verified with the fragment end positions produced by bamtobed tool from BEDTools^29^ to ensure its accuracy.

#### Quality control metrics

CUT&RUNTools reports a number of metrics to evaluate the quality of a CUT&RUN dataset, including: fragment size distribution, adapter content percentage, library size, reads duplication rate, alignment percentage, number of peaks, and enrichment of expected motif. The fragment size is measured by the start and end positions of a pair of reads in paired-end sequencing. Since the experimental protocol enriches short fragments, it is routine to ensure that the fragment size is within the expected range (e.g. <120bp). The quality of sequence reads is evaluated by the adapter content percentage, which is the percentage of reads retained after the read trimming step. For a good quality dataset, the number of reads removed by trimming should be less than 10-15%, mostly corresponding to short fragments. A substantially higher number may indicate technical problems such as self-ligation. The library size, which is defined the number of reads in the sample library, should be at minimum 10 million and ideally at least ~15-20 million. The reads duplication rate is defined as the fraction of paired reads that have identical starts for the first mate and ends for the second mate. A good quality data should typically have a low reads duplication rate (10-15%), although the rate may be higher for factors with an affinity for low complexity regions. The alignment percentage is computed as the percentage of reads that can be mapped concordantly to the reference genome. For a good dataset, the alignment percentage should be high (e.g. >90%). CUT&RUNTools detects peaks by applying MACS2^11^ after filtering out a number of uninteresting regions (including RepeatMasked regions, chromosome M, and any blacklisted regions). In case there is prior knowledge regasrding the exected number of peaks, this may also serve as a guide to evaluate the quality of the data. For transcription factors with known sequence specificity, the enrichment of expected motif should be high at the detected peaks. As there is no single score that captures the overall quality, the users are encouraged to make their own judgement call by considering the collective information.

#### CUT&RUNTools usage

Installation instructions are provided at https://bitbucket.org/qzhudfci/cutruntools/src/default/. To use the pipeline, users first create a new job which entails modifying the provided JSON configuration file with information about the sample fastq file path, output path, SLURM resource requirements, and various settings. Then execute ./create_scripts.py config.json to create a working directory and a set of tailored SLURM submission scripts. Finally, to start the analysis for a sample of interest, users simply execute ./integrated.all.steps.sh GATA1_R1_001.fastq.gz. This script will perform the entire analysis pipeline via a 1-command interface. Options are also available for running the steps of the pipeline individually (see manual on the website for details).

### Code availability

CUT&RUNTools is available at: https://bitbucket.org/qzhudfci/cutruntools/src/default/.

## Acknowledgements

We thank Peter Skene and Steven Henikoff for advice on CUT&RUN protocols; Birgit Knoechel of Dana-Farber Molecular Biology Core Facility for DNA sequencing; Harvard Medical School Research Computing for providing the computing resource for sequencing data analysis; members of Stuart Orkin lab meeting, Daniel Bauer, Alan Cantor for useful feedback. This work was supported by the Howard Hughes Medical Institute (HHMI to S.H.O); National Heart, Lung, and Blood Institute (NHLBI) (R01 HL119099 to G.-C.Y; R01 HL032259 to S.H.O); National Human Genome Research Institute (NHGRI) (HG009663 to G.-C.Y).

## Author Contributions

Q.Z., N.L., S.H.O., G.-C.Y: conceived the project, wrote the paper. Q.Z.: implemented CUT&RUNTools. N.L.: performed CUT&RUN experiments. All authors read and approved the final manuscript.

## References

1. Solomon, M. J. & Varshavsky, A. Formaldehyde-mediated DNA-protein crosslinking: a probe for in vivo chromatin structures. Proc. Natl. Acad. Sci. 82, 6470–6474 (1985).

2. Baranello, L., Kouzine, F., Sanford, S. & Levens, D. ChIP bias as a function of cross-linking time. Chromosom. Res. 24, 175–181 (2016).

3. Meyer, C. A. & Liu, X. S. Identifying and mitigating bias in next-generation sequencing methods for chromatin biology. Nat. Rev. Genet. 15, 709–721 (2014).

4. Teytelman, L., Thurtle, D. M., Rine, J. & van Oudenaarden, A. Highly expressed loci are vulnerable to misleading ChIP localization of multiple unrelated proteins. Proc. Natl. Acad. Sci. 110, 18602–18607 (2013).

5. Skene, P. J. & Henikoff, S. An efficient targeted nuclease strategy for high-resolution mapping of DNA binding sites. Elife 1–35 (2016). doi:10.1101/097188

6. Warfield, L. et al. Transcription of Nearly All Yeast RNA Polymerase II-Transcribed Genes Is Dependent on Transcription Factor TFIID. Mol. Cell 68, 118–129.e5 (2017).

7. Liu, N. et al. Direct Promoter Repression by BCL11A Controls the Fetal to Adult Hemoglobin Switch. Cell 173, 430–442.e17 (2018).

8. Roth, T. L. et al. Reprogramming human T cell function and specificity with non-viral genome targeting. Nature 559, 405–409 (2018).

9. Bolger, A. M., Lohse, M. & Usadel, B. Trimmomatic: A flexible trimmer for Illumina sequence data. Bioinformatics 30, 2114–2120 (2014).

10. Langmead Ben & Steven, S. Fast gapped-read alignment with Bowtie 2. Nat. Methods 9, 357–359 (2013).

11. Zhang, Y. et al. Model-based Analysis of ChIP-Seq (MACS). Genome bio 1–9 (2015). doi:10.1186/gb-2008-9-9-r137)

12. Machanick, P. & Bailey, T. L. MEME-ChIP: Motif analysis of large DNA datasets. Bioinformatics 27, 1696–1697 (2011).

13. Neph, S. et al. An expansive human regulatory lexicon encoded in transcription factor footprints. Nature 489, 83–90 (2012).

14. Hesselberth, J. R. et al. Global mapping of protein-DNA interactions in vivo by digital genomic footprinting. Nat. Methods 6, 283–289 (2009).

15. Pique-Regi, R. \emphet al. et al. Accurate inference of transcription factor binding from DNA sequence and chromatin accessibility data. Genome Res. 21, 447–455 (2011).

16. Pevny, L. et al. Erythroid differentiation in chimaeric mice blocked by a targeted mutation in the gene for transcription factor GATA-1. Nature 349, 257–260 (1991).

17. Hasegawa, A. & Shimizu, R. GATA1 Activity Governed by Configurations of cis-Acting Elements. Front. Oncol. 6, (2017).

18. Wilkinson-White, L. et al. Structural basis of simultaneous recruitment of the transcriptional regulators LMO2 and FOG1/ZFPM1 by the transcription factor GATA1. Proc. Natl. Acad. Sci. 108, 14443–14448 (2011).

19. Wadman, I. A. et al. The LIM-only protein Lmo2 is a bridging molecule assembling an erythroid, DNA-binding complex which includes the TAL1, E47, GATA-1 and Ldb1/NLI proteins. EMBO J. 16, 3145–3157 (1997).

20. Kassouf, M. T. et al. Genome-wide identification of TAL1’s functional targets: Insights into its mechanisms of action in primary erythroid cells. Genome Res. 20, 1064–1083 (2010).

21. Varshney, A. et al. Genetic regulatory signatures underlying islet gene expression and type 2 diabetes. Proc. Natl. Acad. Sci. 114, 2301–2306 (2017).

22. Piper, J. et al. Wellington: A novel method for the accurate identification of digital genomic footprints from DNase-seq data. Nucleic Acids Res. 41, (2013).

23. Buenrostro, J. D., Giresi, P. G., Zaba, L. C., Chang, H. Y. & Greenleaf, W. J. Transposition of native chromatin for fast and sensitive epigenomic profiling of open chromatin, DNA-binding proteins and nucleosome position. Nat. Methods 10, 1213–1218 (2013).

24. Yoo, A. B., Jette, M. A. & Grondona, M. SLURM: Simple Linux Utility for Resource Management. 44–60 (2003). doi:10.1007/10968987_3

25. Pimkin, M. et al. Divergent functions of hematopoietic transcription factors in lineage priming and differentiation during erythro-megakaryopoiesis. Genome Res. 24, 1932–1944 (2014).

26. Vierstra, J. & Stamatoyannopoulos, J. A. Genomic footprinting. Nat. Methods 13, 213–221 (2016).

27. Fu, Y., Wu, P. H., Beane, T., Zamore, P. D. & Weng, Z. Elimination of PCR duplicates in RNA-seq and small RNA-seq using unique molecular identifiers. BMC Genomics 19, (2018).

28. Hahne, F. & Ivanek, R. Visualizing genomic data using Gviz and bioconductor. Methods Mol. Biol. 1418, 335–351 (2016).

29. Quinlan, A. R. BEDTools: The Swiss-Army tool for genome feature analysis. Curr. Protoc. Bioinforma. 2014, 11.12.1–11.12.34 (2014).

