## Supplementary Material for "CUT&RUNTools: a flexible pipeline for CUT&RUN processing and footprint analysis"

### Supplementary Fig 1

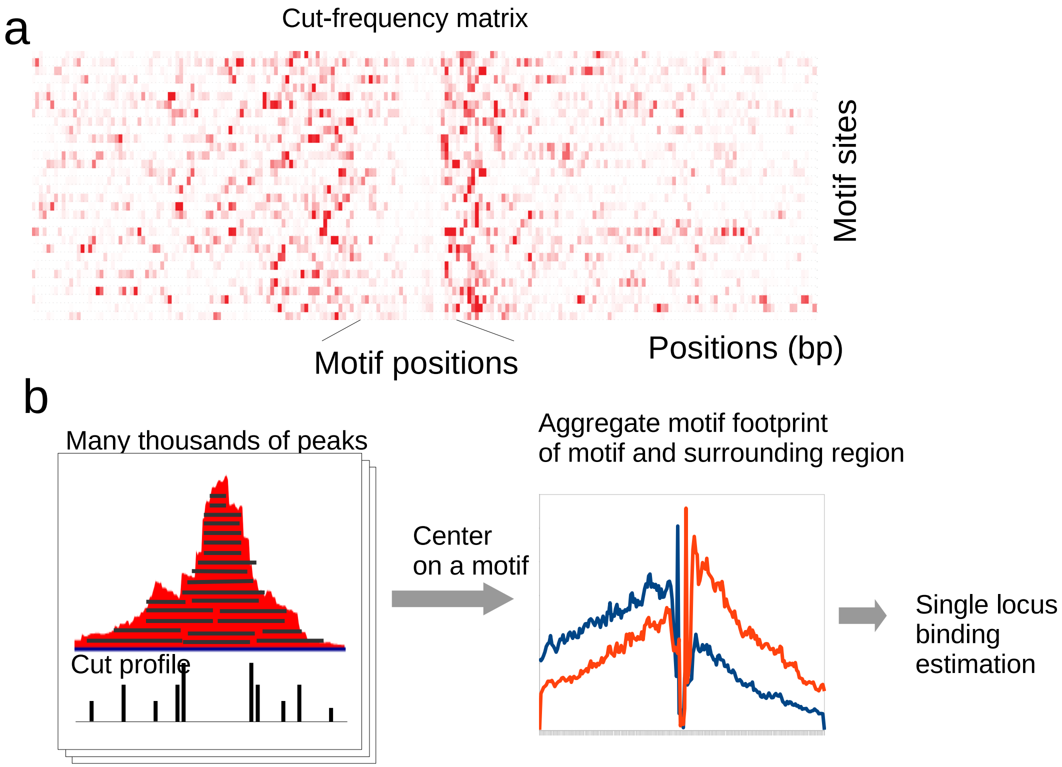

### Supplementary Fig 2

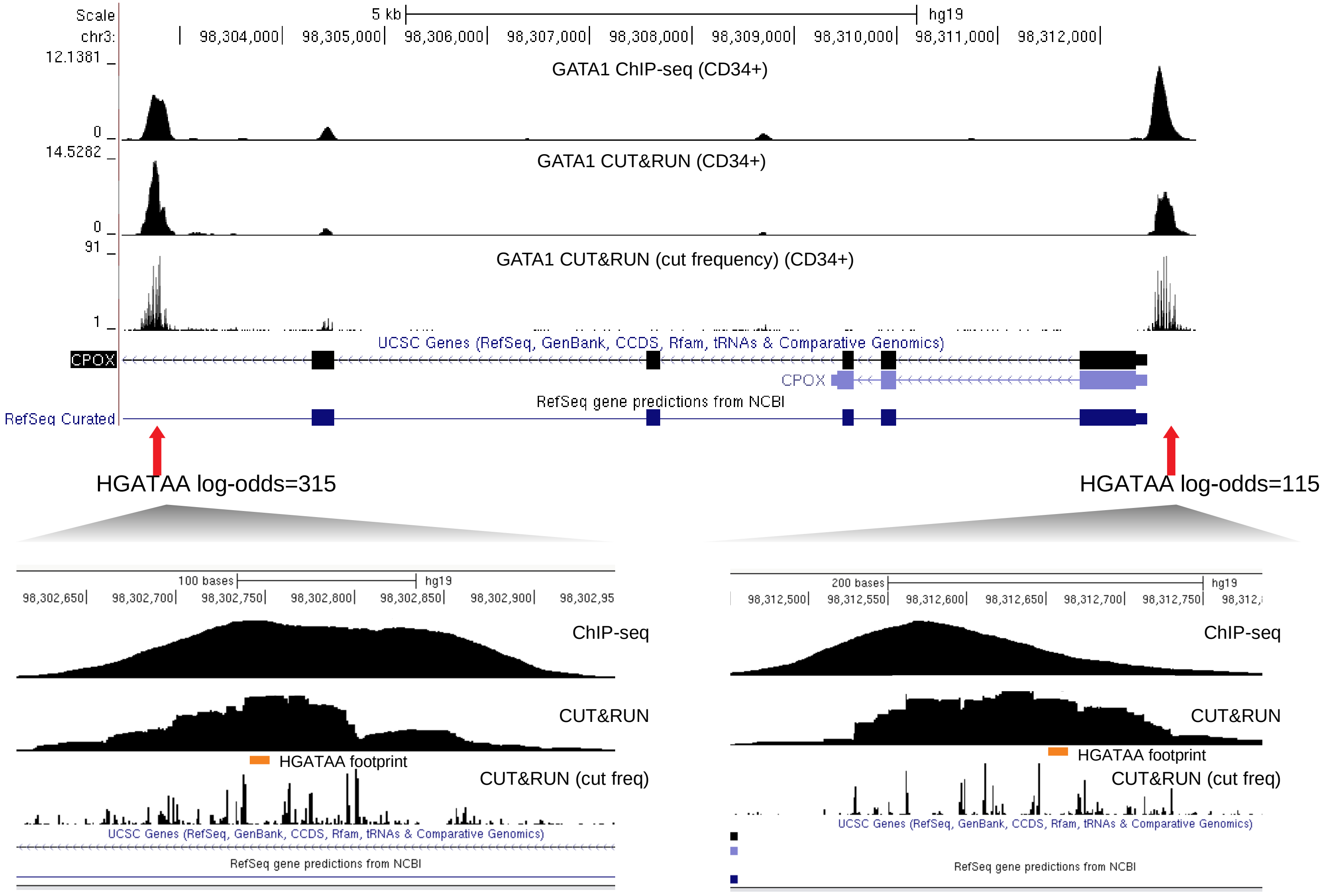

### Supplementary Fig 3

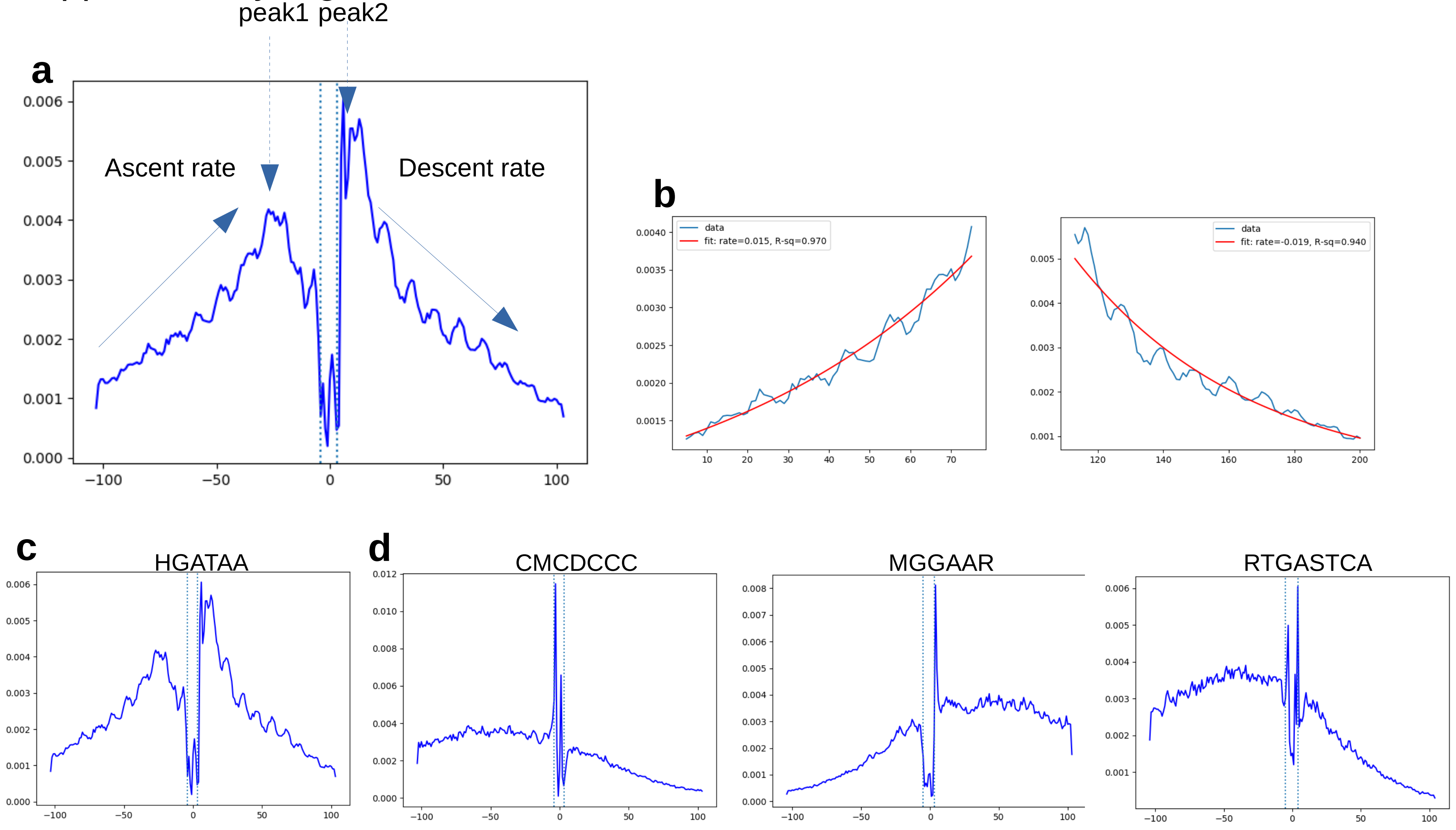

### Supplementary Fig 4

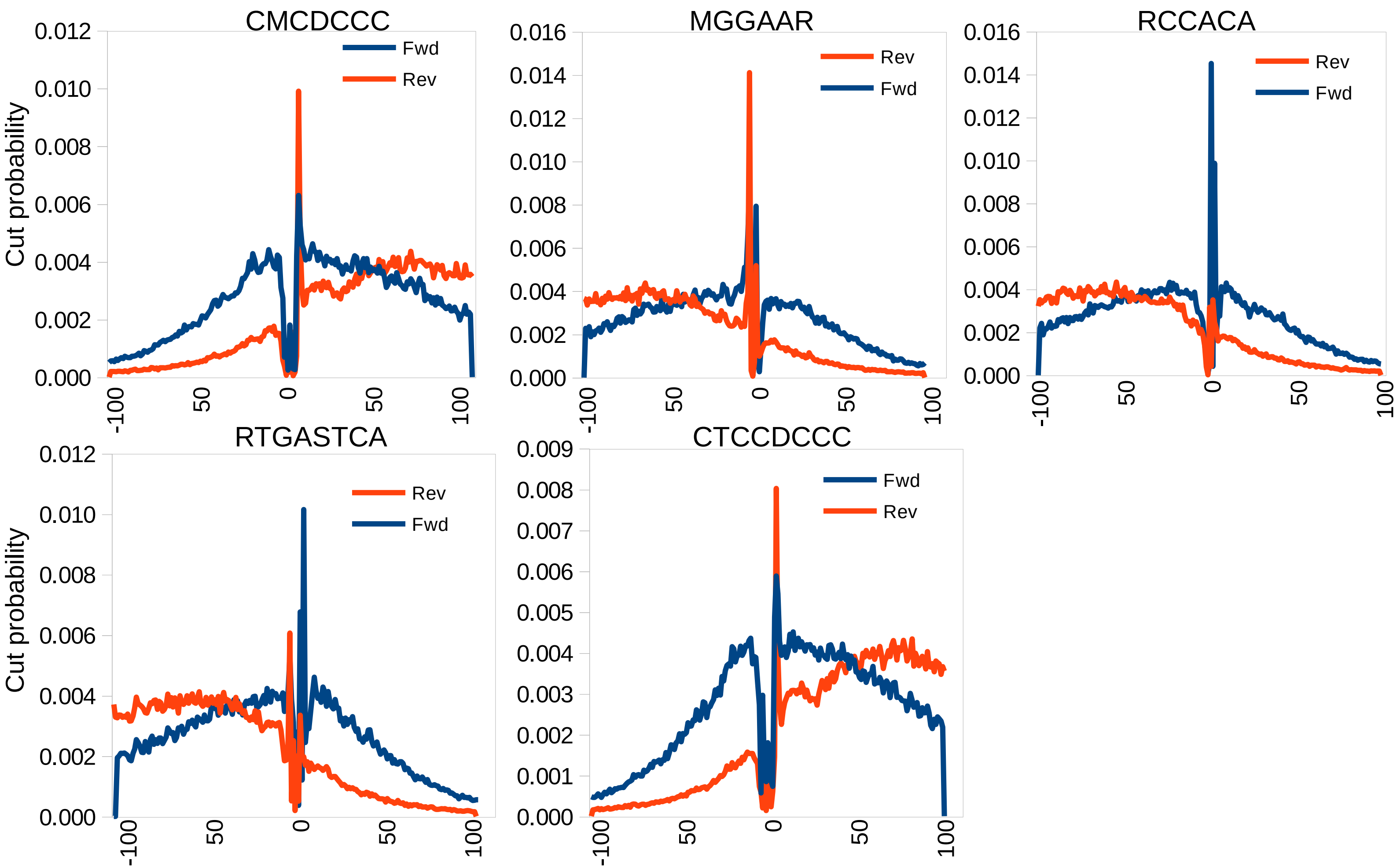

### Supplementary Fig 5

#### GATA1-TAL1 composite

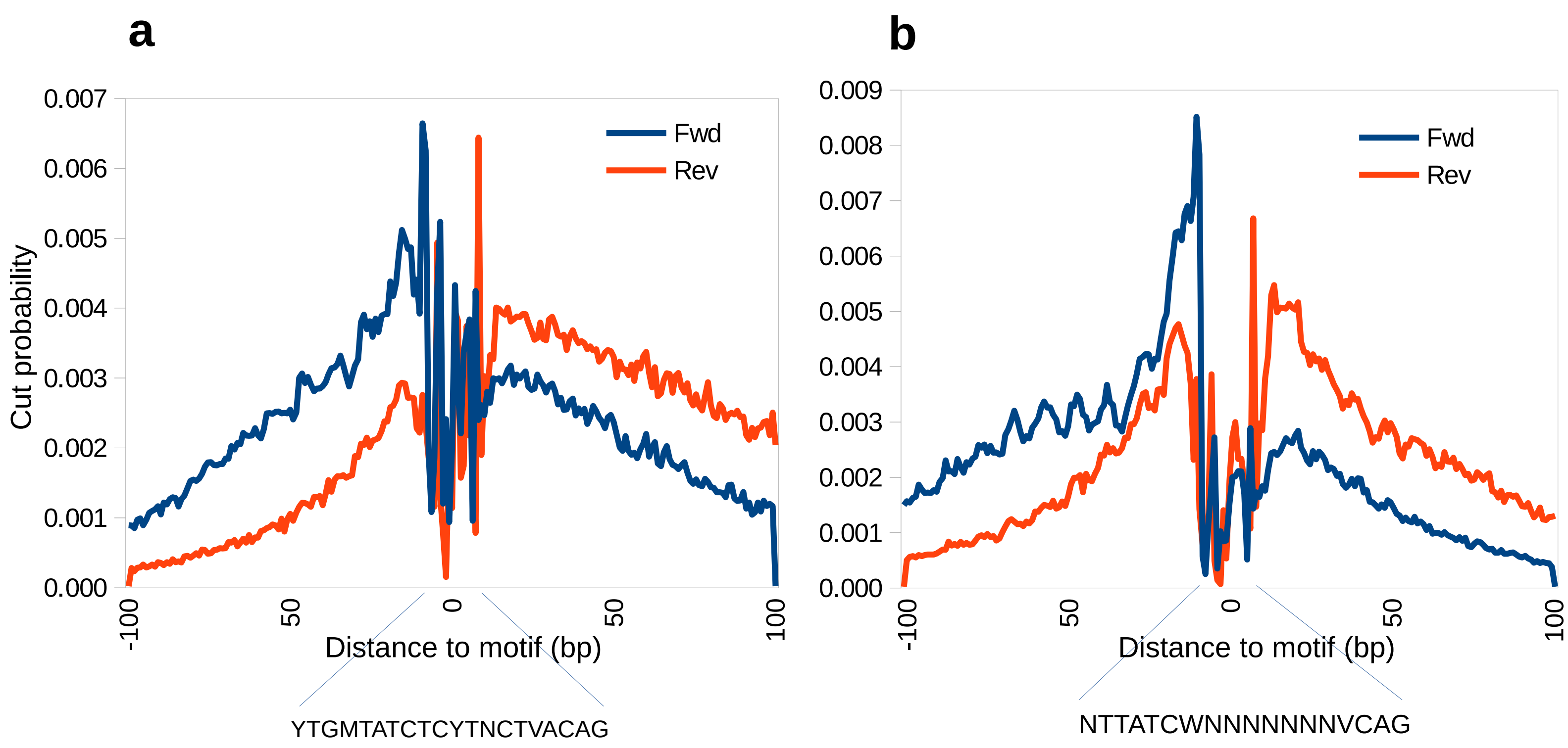

### Supplementary Fig 6

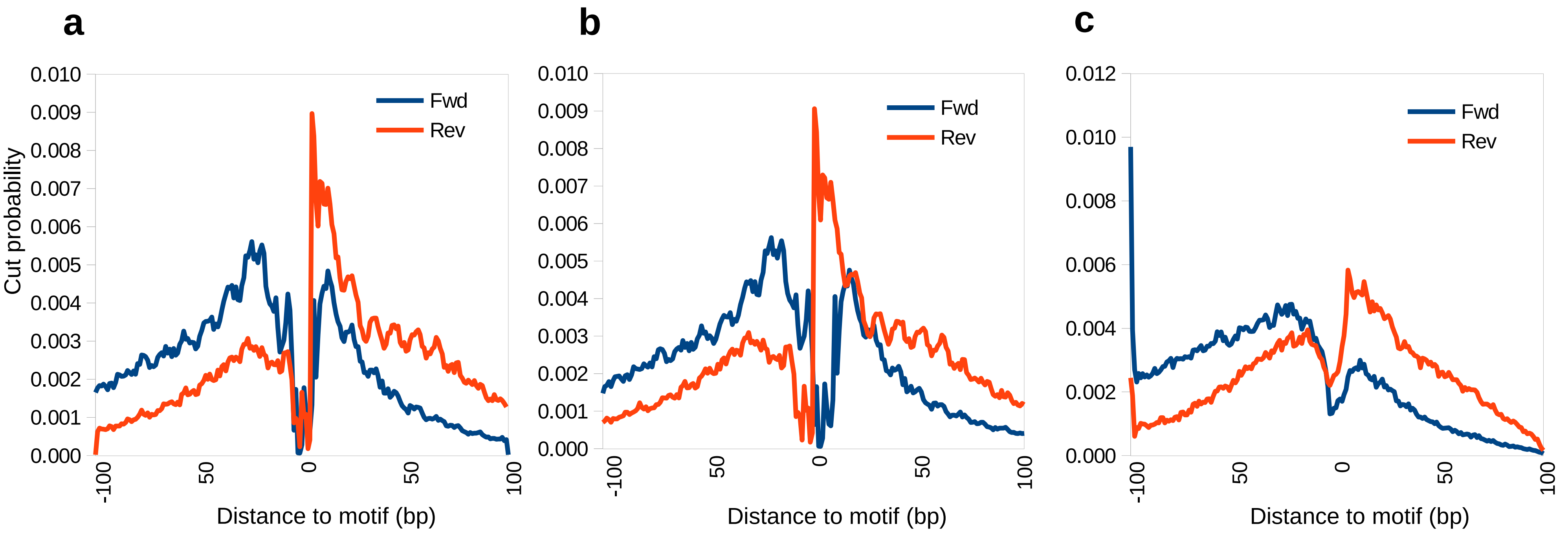

### Supplementary Fig 7

a

Chr1:32510-32516. HGATAA on (-) strand

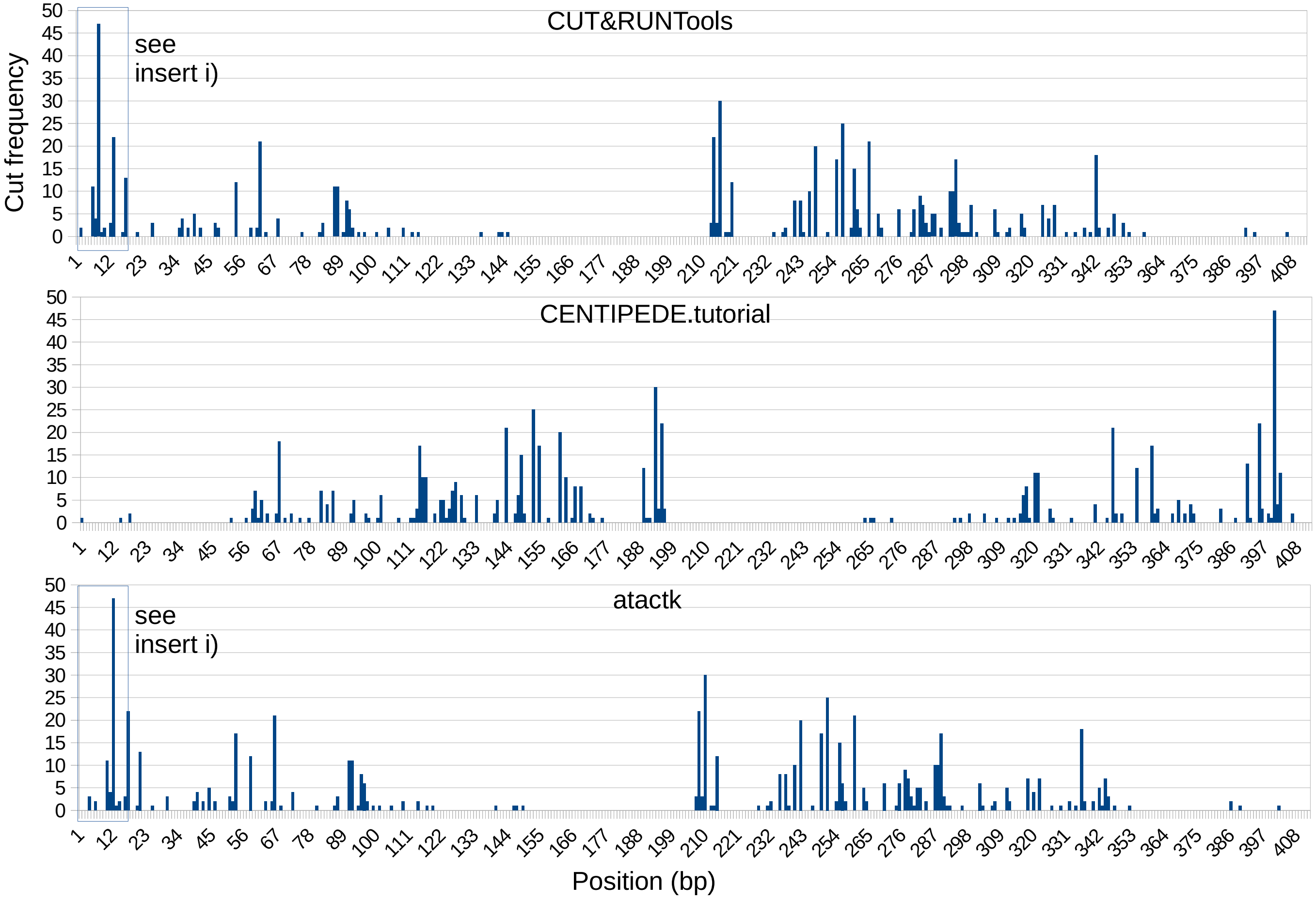


Correct reference.

Incorrect. Reason: Cut frequency should be reversed for this site since the motif is located on (-) strand.

Incorrect. Reason: Cut frequency should be shifted by 4nt. ATAC-seq transpose leaves 4nt overhang which does not apply in CUTRUN.

i

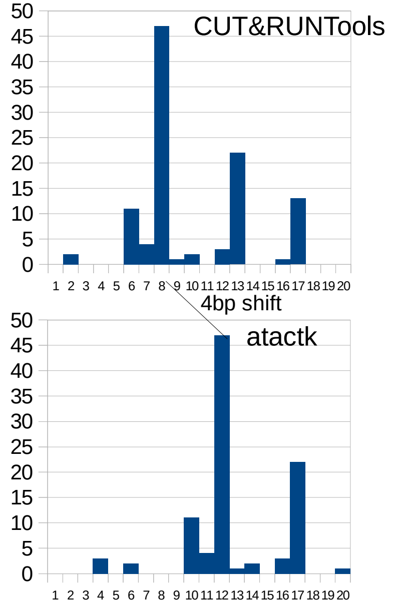

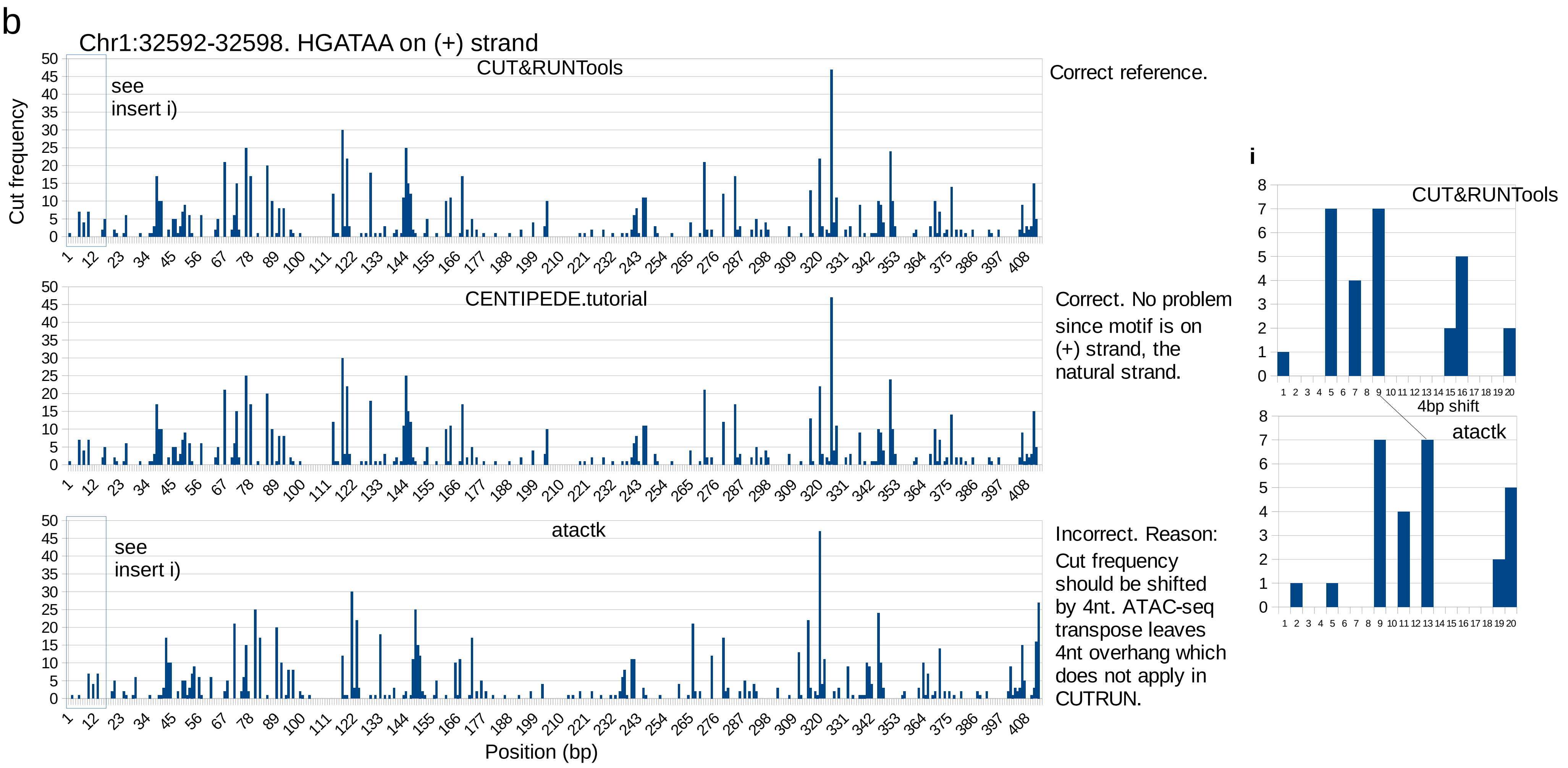
b
Chr1:32592-32598. HGATAA on (+) strand
CUT&RUNTools
see insert i)
Correct reference.
CENTIPEDE.tutorial
Correct. No problem since motif is on (+) strand, the natural strand.
atactk
see insert i)
Incorrect. Reason: Cut frequency should be shifted by 4nt. ATAC-seq transpose leaves 4nt overhang which does not apply in CUTRUN.
Cut frequency
Position (bp)
i
CUT&RUNTools
4bp shift
atactk

### Supplementary Fig 8

#### Anatomy of DNA fragment

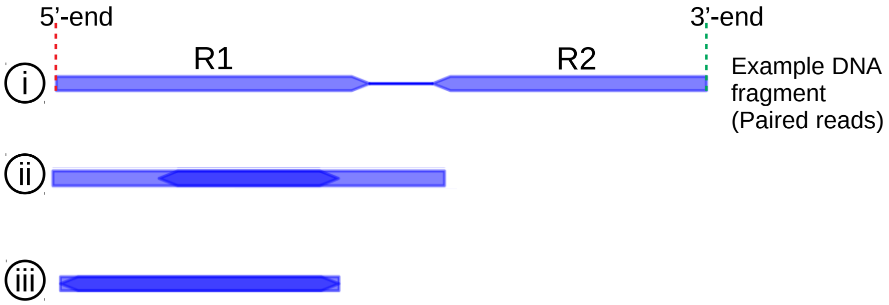

### Supplementary Fig 9

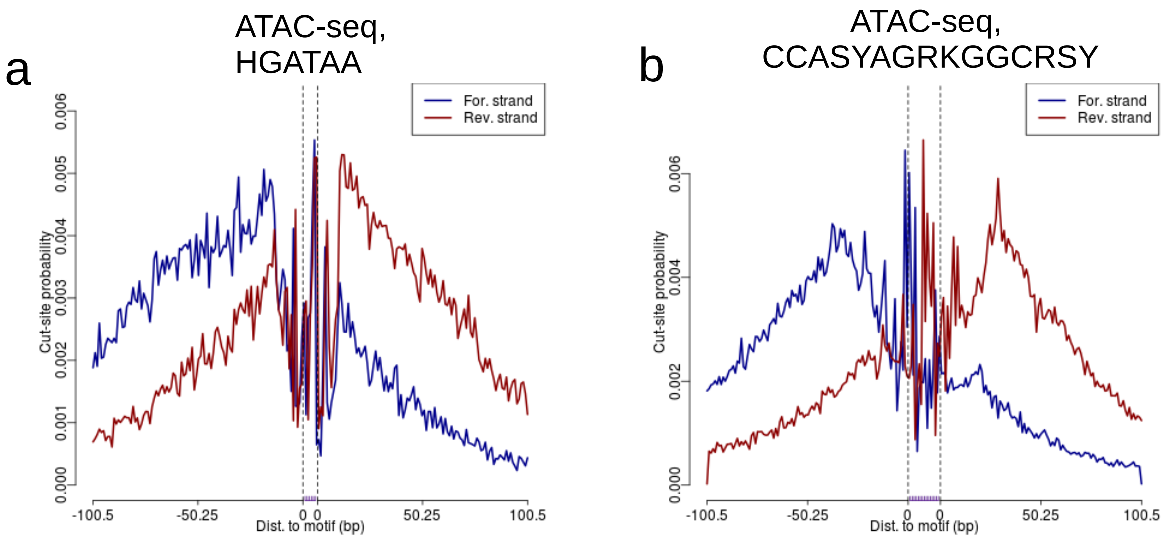

### Supplementary Fig 10

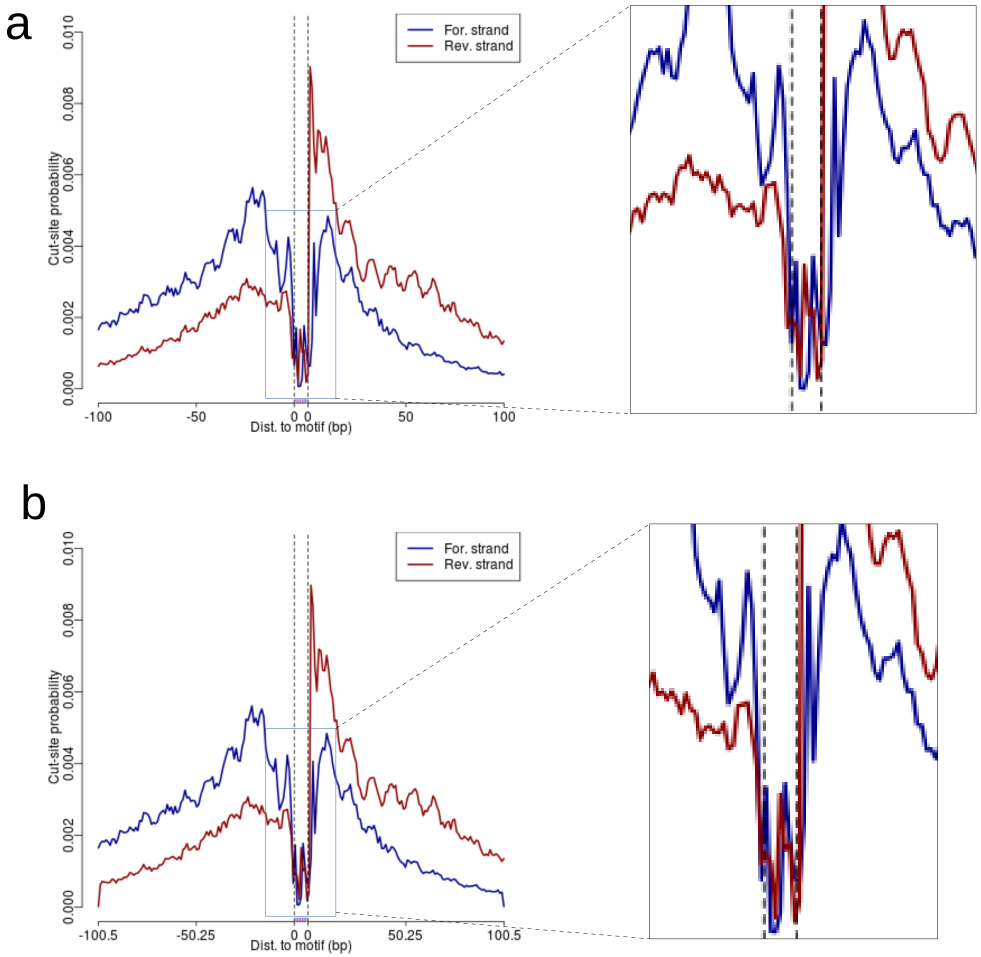

Supplementary Table 1: improvement of reads utilization by custom trimming, dovetail alignment

| | Setting | Number of reads | Aligned concordantly 1 or more times | | Total | Alignment rate |
|---|---|---|---|---|---|---|
| | | | 1 time | >1 times | | |
| CUT&RUNTools | default | 28787551 | 13070514 | 15380579 | 28451093 | 0.9883 |
| | no dovetail | 28787551 | 12804553 | 15075315 | 27879868 | 0.9685 |
| | no trimming | 31515085 | 11288822 | 12932597 | 24221419 | 0.7686 |
| | no trimming and no dovetail | 31515085 | 9003955 | 11327689 | 20331644 | 0.6451 |

### Supplementary Table 2: Binding log-odds* for HGATAA motif from GATA1 CUT&RUN

| Chromosome | Start | End | Motif | - | Orientation | Log-odds |
|---|---|---|---|---|---|---|
| chr1 | 206084286 | 206084292 | 1-HGATAA-1-chr1 | 31.5 | - | 539.94350133 |
| chr3 | 128775880 | 128775886 | 1-HGATAA-1-chr3 | 34.5 | + | 539.53606379 |
| chr19 | 51161579 | 51161585 | 1-HGATAA-2-chr19 | 34.5 | + | 538.99925407 |
| chr15 | 66097635 | 66097641 | 1-HGATAA-3-chr15 | 31.5 | + | 538.70591937 |
| chr15 | 66097669 | 66097675 | 1-HGATAA-1-chr15 | 34.5 | - | 526.8836613 |
| chr21 | 46831592 | 46831598 | 1-HGATAA-3-chr21 | 30 | - | 511.70845847 |
| chr9 | 135674727 | 135674733 | 1-HGATAA-1-chr9 | 34.5 | + | 488.71658446 |
| chr14 | 103844534 | 103844540 | 1-HGATAA-1-chr14 | 34.5 | - | 483.96802505 |
| chr4 | 154001237 | 154001243 | 1-HGATAA-1-chr4 | 34.5 | - | 445.57355961 |
| chr17 | 40781892 | 40781898 | 1-HGATAA-1-chr17 | 34.5 | + | 421.86582392 |
| chr8 | 21721189 | 21721195 | 1-HGATAA-2-chr8 | 31.5 | + | 407.023727 |
| chr20 | 4173122 | 4173128 | 1-HGATAA-2-chr20 | 31.5 | + | 404.41183257 |
| chr15 | 31270710 | 31270716 | 1-HGATAA-2-chr15 | 34.5 | + | 402.57658532 |
| chr4 | 144924560 | 144924566 | 1-HGATAA-2-chr4 | 31.5 | - | 401.80075706 |
| chr12 | 122309495 | 122309501 | 1-HGATAA-2-chr12 | 31.5 | - | 396.81915852 |
| chr17 | 31224658 | 31224664 | 1-HGATAA-1-chr17 | 34.5 | + | 386.97191045 |
| chr4 | 144806254 | 144806260 | 1-HGATAA-2-chr4 | 31.5 | - | 377.69801447 |
| chr17 | 17155717 | 17155723 | 1-HGATAA-2-chr17 | 34.5 | + | 369.99541714 |
| chr3 | 42878291 | 42878297 | 1-HGATAA-2-chr3 | 34.5 | + | 369.53921139 |
| chr14 | 31529077 | 31529083 | 1-HGATAA-2-chr14 | 34.5 | + | 359.84514632 |
| chr6 | 4177295 | 4177301 | 1-HGATAA-1-chr6 | 34.5 | - | 352.63599767 |
| chr3 | 158461534 | 158461540 | 1-HGATAA-2-chr3 | 31.5 | - | 352.23946823 |
| chr14 | 23366729 | 23366735 | 1-HGATAA-1-chr14 | 34.5 | + | 348.54592287 |
| chr2 | 182170724 | 182170730 | 1-HGATAA-2-chr2 | 31.5 | + | 347.54053257 |
| chr8 | 131355530 | 131355536 | 1-HGATAA-1-chr8 | 31.5 | + | 346.2938888 |
| chr11 | 12222906 | 12222912 | 1-HGATAA-2-chr11 | 31.5 | + | 344.54880589 |
| chr12 | 47601134 | 47601140 | 1-HGATAA-1-chr12 | 34.5 | + | 344.39714697 |
| chr2 | 106002546 | 106002552 | 1-HGATAA-1-chr2 | 34.5 | + | 331.66061495 |
| chr1 | 179016337 | 179016343 | 1-HGATAA-1-chr1 | 34.5 | + | 327.47086225 |
| chr19 | 51161526 | 51161532 | 1-HGATAA-1-chr19 | 34.5 | + | 326.16855012 |
| chr11 | 10764567 | 10764573 | 1-HGATAA-2-chr11 | 31.5 | + | 324.70144216 |
| chr4 | 154001272 | 154001278 | 1-HGATAA-2-chr4 | 31.5 | - | 324.58530837 |
| chr17 | 35085016 | 35085022 | 1-HGATAA-3-chr17 | 34.5 | + | 323.51680601 |
| chr11 | 5297187 | 5297193 | 1-HGATAA-1-chr11 | 34.5 | + | 322.13082367 |
| chr6 | 10750100 | 10750106 | 1-HGATAA-1-chr6 | 34.5 | - | 321.63525671 |
| chr16 | 21548326 | 21548332 | 1-HGATAA-2-chr16 | 31.5 | + | 319.85751909 |
| chr12 | 116635110 | 116635116 | 1-HGATAA-1-chr12 | 34.5 | - | 318.83571936 |
| chr14 | 65510005 | 65510011 | 1-HGATAA-1-chr14 | 34.5 | + | 318.63418452 |
| chr1 | 114457022 | 114457028 | 1-HGATAA-1-chr1 | 34.5 | + | 318.49413784 |
| chr3 | 98302746 | 98302752 | 1-HGATAA-1-chr3 | 34.5 | + | 315.45546771 |
| chr10 | 30726160 | 30726166 | 1-HGATAA-1-chr10 | 34.5 | + | 311.750006 |
| chr3 | 38765774 | 38765780 | 1-HGATAA-3-chr3 | 31.5 | + | 311.39122098 |
| chr2 | 7162991 | 7162997 | 1-HGATAA-2-chr2 | 31.5 | - | 308.06039247 |
| chr11 | 72767188 | 72767194 | 1-HGATAA-1-chr11 | 34.5 | - | 307.06930345 |
| chr12 | 122309544 | 122309550 | 1-HGATAA-1-chr12 | 34.5 | - | 302.48306998 |
| chr1 | 160959833 | 160959839 | 1-HGATAA-1-chr1 | 34.5 | - | 301.71039986 |
| chr3 | 46550819 | 46550825 | 1-HGATAA-1-chr3 | 34.5 | + | 298.31930536 |
| chr6 | 42060095 | 42060101 | 1-HGATAA-1-chr6 | 34.5 | + | 297.00240629 |
| chr2 | 69828488 | 69828494 | 1-HGATAA-1-chr2 | 34.5 | + | 296.53737507 |
| chr10 | 103210225 | 103210231 | 1-HGATAA-2-chr10 | 31.5 | - | 293.99385537 |

* Top 50 sites with highest binding log odds are shown.

#### Supplementary Table 3: Footprint shape and symmetry analysis* (GATA1 CUT&RUN)

| Motif | Ascent rate (1) | $R^2$ (coefficient determination) | Peak1 position | Descent rate (2) | $R^2$ (coefficient determination) | Peak2 position | Footprint Symmetry Score | Δrates (1-2) |
|---|---|---|---|---|---|---|---|---|
| DREME-16.BCTTATC | 0.017885 | 0.952077 | 91 | -0.015968 | 0.972970 | 129 | 0.032564 | 0.001917 |
| DREME-1.HGATAA | 0.014891 | 0.969814 | 75 | -0.018961 | 0.940017 | 113 | 0.032266 | 0.004070 |
| MEME-1.BBCTTATCTBH | 0.018207 | 0.941735 | 92 | -0.015034 | 0.972119 | 130 | 0.031761 | 0.003173 |
| DREME-14.CTGATTRG | 0.022813 | 0.982532 | 77 | -0.007001 | 0.929999 | 114 | 0.028925 | 0.015812 |
| MEME-50.GGATAAGCACC | 0.007829 | 0.937589 | 80 | -0.021961 | 0.967465 | 113 | 0.028587 | 0.014132 |
| DREME-7.AGATTA | 0.006539 | 0.801149 | 76 | -0.023579 | 0.967788 | 113 | 0.028058 | 0.017040 |
| DREME-12.CTGATAKS | 0.008038 | 0.932221 | 76 | -0.020616 | 0.984048 | 110 | 0.027780 | 0.012578 |
| MEME-5.TYATTCRKCTCRGSWWGRTGASTCAGRGCC | 0.006759 | 0.887717 | 75 | -0.021965 | 0.979954 | 132 | 0.027525 | 0.015206 |
| DREME-19.AAAAAHAA | 0.025478 | 0.987155 | 93 | -0.004194 | 0.515339 | 173 | 0.027312 | 0.021285 |
| DREME-9.CTCCDCCC | 0.024397 | 0.990011 | 90 | -0.003625 | 0.693915 | 145 | 0.026668 | 0.020771 |
| DREME-6.RTGASTCA | 0.005535 | 0.836229 | 62 | -0.022152 | 0.984747 | 113 | 0.026443 | 0.016618 |
| MEME-8.CGGCCCCGC | 0.023271 | 0.983471 | 93 | -0.004910 | 0.705149 | 147 | 0.026348 | 0.018361 |
| MEME-37.TCCTGCTSTKG | 0.023555 | 0.979295 | 91 | -0.003879 | 0.823788 | 139 | 0.026263 | 0.019676 |
| MEME-32.RCTGCCMTCTYVTGS | 0.021665 | 0.973954 | 93 | -0.005579 | 0.898509 | 142 | 0.026114 | 0.016086 |
| MEME-20.GCTRYSAGTGABAGAGAMCA | 0.005315 | 0.874646 | 78 | -0.021758 | 0.979390 | 122 | 0.025958 | 0.016442 |
| MEME-33.CCCAGGCGTGG | 0.005168 | 0.805398 | 59 | -0.022106 | 0.984650 | 114 | 0.025930 | 0.016938 |
| DREME-2.CMCDCCC | 0.022766 | 0.984864 | 90 | -0.004861 | 0.719514 | 145 | 0.025919 | 0.017905 |
| DREME-8.CAGMWG | 0.004109 | 0.834260 | 74 | -0.022751 | 0.975606 | 112 | 0.025624 | 0.018642 |
| MEME-29.TWKCWRNMASCCRSCACASAG | 0.005768 | 0.905974 | 83 | -0.020632 | 0.981285 | 124 | 0.025472 | 0.014864 |
| MEME-30.TMTATCTSTGTDCTSCTTGKC | 0.006531 | 0.922546 | 89 | -0.019674 | 0.972762 | 122 | 0.025163 | 0.013143 |
| MEME-3.GAKGKRRTCAGASNCTGGRCTGAGWGAAG | 0.004464 | 0.791282 | 95 | -0.021919 | 0.984111 | 132 | 0.025103 | 0.017455 |
| MEME-15.GCWCTGCCTCC | 0.003754 | 0.740657 | 65 | -0.024095 | 0.913687 | 112 | 0.024796 | 0.020341 |
| DREME-17.CCWCCTCC | 0.024659 | 0.990057 | 88 | -0.001191 | 0.181527 | 110 | 0.024630 | 0.023469 |
| MEME-47.CAGCCCCACCC | 0.023026 | 0.985668 | 93 | -0.002392 | 0.518524 | 120 | 0.023936 | 0.020634 |
| DREME-10.CHGCC | 0.004036 | 0.700540 | 70 | -0.021109 | 0.982761 | 109 | 0.023572 | 0.017073 |
| MEME-35.CATCWCAGCCA | 0.003658 | 0.715174 | 78 | -0.021315 | 0.982304 | 112 | 0.023554 | 0.017657 |
| MEME-2.GCCCCGCCCTC | 0.022955 | 0.988390 | 95 | -0.002275 | 0.327758 | 110 | 0.023434 | 0.020680 |
| DREME-11.AMACAS | 0.004115 | 0.711195 | 55 | -0.021011 | 0.974483 | 111 | 0.023402 | 0.016897 |
| DREME-18.CWGTSAC | 0.004215 | 0.750771 | 64 | -0.020574 | 0.963616 | 109 | 0.022989 | 0.016359 |
| DREME-5.RCCACA | 0.003010 | 0.662202 | 79 | -0.020842 | 0.982252 | 110 | 0.022465 | 0.017831 |
| DREME-20.AGGCGTGK | 0.020585 | 0.978716 | 96 | -0.003452 | 0.656984 | 126 | 0.022415 | 0.017133 |
| DREME-4.MGGAAR | 0.001027 | 0.116794 | 96 | -0.020259 | 0.967402 | 113 | 0.019719 | 0.019232 |
| MEME-34.GGVCMCAGAGG | 0.001975 | 0.280822 | 96 | -0.019529 | 0.955259 | 114 | 0.019210 | 0.017554 |

* For each footprint, we fit the data with A exp(B * x). Ascent rate refers to the parameter B estimated on the left arm of the footprint. Descent rate refers to the parameter B estimated on the right arm of the footprint. See **Supplementary Fig 3**.

Supplementary Data 1: Summary of motifs found for GATA1 CUT&RUN (based on a subset of 5000 peaks)

| MOTIF_INDEX | MOTIF_SOURCE | MOTIF_ID | FACTOR | ALT_ID | WIDTH | SITES | E-VALUE |
|---|---|---|---|---|---|---|---|
| 1 | DREME | HGATAA | GATA1 | DREME-1 | 6 | 7321 | 1.1e-1163 |
| 2 | MEME | BBCTTATCTBH | GATA1 | MEME-1 | 11 | 344 | 2.6e-1082 |
| 3 | DREME | AGATA | GATA1 | DREME-3 | 5 | 2771 | 6.9e-688 |
| 4 | DREME | BCTTATC | GATA1 | DREME-16 | 7 | 176 | 7.9e-676 |
| 5 | DREME | CMCDCCC | KLF1 | DREME-2 | 7 | 2014 | 5.9E-94 |
| 6 | DREME | AGATTA | | DREME-7 | 6 | 1064 | 1.1E-86 |
| 7 | DREME | CTGATAKS | GATA1 | DREME-12 | 8 | 250 | 5.2E-58 |
| 8 | DREME | MGGAAR | ETS1, FLI1 | DREME-4 | 6 | 2923 | 1.5E-41 |
| 9 | DREME | CAGMWG | | DREME-8 | 6 | 3901 | 2.7E-40 |
| 10 | DREME | RTGASTCA | NFE2 | DREME-6 | 8 | 433 | 6.3E-35 |
| 11 | DREME | RCCACA | RUNX1 | DREME-5 | 6 | 1706 | 4.5E-33 |
| 12 | DREME | CHGCC | | DREME-10 | 5 | 6795 | 3E-29 |
| 13 | DREME | CTGATTRG | | DREME-14 | 8 | 173 | 2.4E-26 |
| 14 | MEME | GGATAAGCACC | | MEME-50 | 11 | 4 | 1.30E-24 |
| 15 | MEME | YTGMTATCTCYTNCTVACAG | GATA1/TAL1 | MEME-12 | 20 | 20 | 5.80E-24 |
| 16 | DREME | CTCCDCCC | KLF1 | DREME-9 | 8 | 411 | 3E-21 |
| 17 | DREME | AMACAS | | DREME-11 | 6 | 2359 | 4.4E-16 |
| 18 | DREME | CWGTSAC | PBX3/MEIS | DREME-18 | 7 | 512 | 4.3E-13 |
| 19 | MEME | TMTATCTSTGTDCTSCTTGKC | | MEME-30 | 21 | 8 | 1.8E-11 |
| 20 | DREME | RGAAA | | DREME-13 | 5 | 4901 | 8.5E-11 |
| 21 | MEME | RCTGCCMTCTYVTGS | | MEME-32 | 15 | 13 | 9.1E-11 |
| 22 | MEME | GCCCCGCCCTC | | MEME-2 | 11 | 36 | 3.1E-08 |
| 23 | DREME | ACGT | | DREME-15 | 4 | 909 | 5.9E-08 |
| 24 | MEME | CATCWCAGCCA | GFI1B | MEME-35 | 11 | 11 | 6.4E-06 |
| 25 | MEME | GCTRYSAGTGABAGAGAMCA | ZBTB3 | MEME-20 | 20 | 10 | 1.7E-05 |
| 26 | DREME | CCWCCTCC | | DREME-17 | 8 | 154 | 1.9E-05 |
| 27 | MEME | TWKCWRNMASCCRSCACASAG | | MEME-29 | 21 | 11 | 3.3E-05 |
| 28 | MEME | GGVCMCAGAGG | | MEME-34 | 11 | 9 | 7.5E-05 |
| 29 | MEME | TYATTCRKCTCRGSWWGRTGASTCAGRGCC | | MEME-5 | 30 | 3 | 0.00017 |
| 30 | DREME | AGGCGTGK | | DREME-20 | 8 | 52 | 0.00052 |
| 31 | MEME | CCCAGGCGTGG | | MEME-33 | 11 | 2 | 0.0011 |
| 32 | DREME | AAAAAHAA | | DREME-19 | 8 | 240 | 0.0016 |
| 33 | MEME | CAGCCCCACCC | | MEME-47 | 11 | 2 | 0.0022 |
| 34 | MEME | CGGCCCCGC | | MEME-8 | 9 | 2 | 0.013 |
| 35 | MEME | GAKGKRRTCAGASNCTGGRCTGAGWGAAG | | MEME-3 | 29 | 7 | 0.02 |
| 36 | MEME | TCCTGCTSTKG | ZIC1/2 | MEME-37 | 11 | 11 | 0.026 |
| 37 | MEME | GCWCTGCCTCC | | MEME-15 | 11 | 37 | 0.029 |
